# Does Hemispheric Asymmetry Reduction in Older Adults (HAROLD) in motor cortex reflect compensation?

**DOI:** 10.1101/2021.06.02.446015

**Authors:** Ethan Knights, Alexa Morcom, Cam-CAN, Richard N. Henson

## Abstract

Older adults tend to display greater brain activation in the non-dominant hemisphere during even basic sensorimotor responses. It is debated whether this Hemispheric Asymmetry Reduction in Older Adults (HAROLD) reflects a compensatory mechanism. Across two independent fMRI experiments involving an adult-lifespan human sample (N = 586 and N = 81; approximately half female) who performed right hand finger responses, we distinguished between these hypotheses using behavioural and multivariate Bayes (MVB) decoding approaches. Standard univariate analyses replicated a HAROLD pattern in motor cortex, but in- and out-of-scanner behavioural results both demonstrated evidence against a compensatory relationship, in that reaction time measures of task performance in older adults did not relate to ipsilateral motor activity. Likewise, MVB showed that this increased ipsilateral activity in older adults did not carry additional information, and if anything, combining ipsilateral with contralateral activity patterns reduced action decoding in older adults (at least in Experiment 1). These results contradict the hypothesis that HAROLD is compensatory, and instead suggest that the age-related, ipsilateral hyper-activation is non-specific, in line with alternative hypotheses about age-related reductions in neural efficiency/differentiation or inter-hemispheric inhibition.

**Significance Statement:** A key goal in the cognitive neuroscience of ageing is to provide a mechanistic explanation of how brain-behaviour relationships change with age. One interpretation of the common finding that task-based hemispheric activity becomes more symmetrical in older adults, is that this shift reflects a compensatory mechanism, with the non-dominant hemisphere needing to “help out” with computations normally performed by the dominant hemisphere. Contrary to this view, our behavioural and brain data indicate that the additional activity in ipsilateral motor cortex in older adults is not reflective of better task performance nor better neural representations of finger actions.

## Introduction

Functional neuroimaging has established that increased age is linked to weaker task-based neural lateralisation (e.g., Cabeza et al., 1997), with older adults showing increased activation of the non-dominant hemisphere; a pattern summarised as Hemispheric Asymmetry Reduction in Older Adults, or “HAROLD” (Cabeza, 2002). The explanation for this reduced lateralisation is debated. A widely cited idea is that the recruitment of the non-dominant hemisphere reflects compensatory mechanisms (Cabeza et al., 2018). An alternative hypothesis is that this increased activation is non-functional (e.g., Grady et al. 1994), perhaps reflecting inefficient or more dedifferentiated neural processing (Morcom & Johnson, 2015).

Motor responses, such as finger (Mattay et al., 2002; Rowe et al., 2006), wrist (Heuninckx et al., 2005) or grasping (Ward & Frackowiak, 2003; Ward et al., 2008) movements, are sufficient to evoke HAROLD patterns in motor areas. For example, mean activation within the right (ipsilateral) motor cortex increases with age when participants respond with their right hand (Tsvetanov et al., 2015). Brain-behaviour relationships are commonly examined to adjudicate between the compensation and inefficiency hypotheses. If ipsilateral activity is compensatory, averaged activation will be positively related to behavioural performance. Nevertheless, “univariate” activation results are inconclusive: greater ipsilateral motor activation in older adults has been reported to show positive (Mattay et al., 2002; Heuninckx et al., 2008), negative (Langan, et al., 2010; Cassady et al., 2020), or no (Riecker et al., 2006) relationship with kinematics. Multivariate approaches offer an alternative way to test these competing hypotheses. If increased ipsilateral activity is compensatory (rather than non-functional), it should contain task-relevant information.

Multivoxel pattern analysis (MVPA) has demonstrated that, in line with de-differentiation, the distinctiveness of information represented within ipsilateral motor areas during finger tapping is reduced in older adults (Carp et al., 2011). However, a stronger assessment of whether ipsilateral motor activity is compensatory requires testing whether task-relevant information in ipsilateral cortex is complementary to that in contralateral cortex. The degree of complementarity could increase with age, even if the total amount of information in ipsilateral cortex decreases with age, as Carp et al. found (i.e., the greater information in young people in ipsilateral cortex could be redundant with that in contralateral cortex). This can be tested by combining voxels across hemispheres and testing whether decoding is improved relative to using voxels from the contralateral hemisphere alone (Morcom & Henson, 2018).

Morcom and Henson (2018) used multivariate Bayes (MVB), a model-based MVPA technique, to test whether one model (set of voxels) is more likely than another in predicting experimental conditions (Friston et al., 2008; Morcom & Friston, 2012). They tested a different ageing-related hypothesis (the Posterior-to-Anterior Shift with Age), which claims that increased anterior activity in older people is also compensatory (Davis et al., 2008). They found that, when predicting memory, Bayesian Model Evidence in older adults was more often reduced, rather than increased, for a model with voxels from both anterior and posterior brain regions compared to a model with posterior voxels only. That is, results were more consistent with the hypothesis that age reduces the efficiency/differentiation of neural activity, rather than compensation.

Here, we applied the same MVB logic to test HAROLD in the context of motor activity related to simple finger presses across two motor fMRI experiments in the “Cam-CAN” population-derived adult lifespan sample (www.cam-can.org; Shafto et al., 2014). In Experiment 1, participants (N=586) pressed a button with their right index finger whenever they saw/heard a visual/auditory stimulus. In Experiment 2, participants (N=81) were cued to press the button under one of four fingers of their right hand (Figure 1). First, we assessed whether greater mean ipsilateral sensorimotor cortex activation was associated with improved (i.e., shorter/less variable) reaction times (RTs) for older adults during the scanner task, and in separate tasks run outside the scanner. Second, we used MVB to test whether the model evidence based on action decoding was “boosted” for older adults when models included ipsilateral voxels.

**Figure 1.**
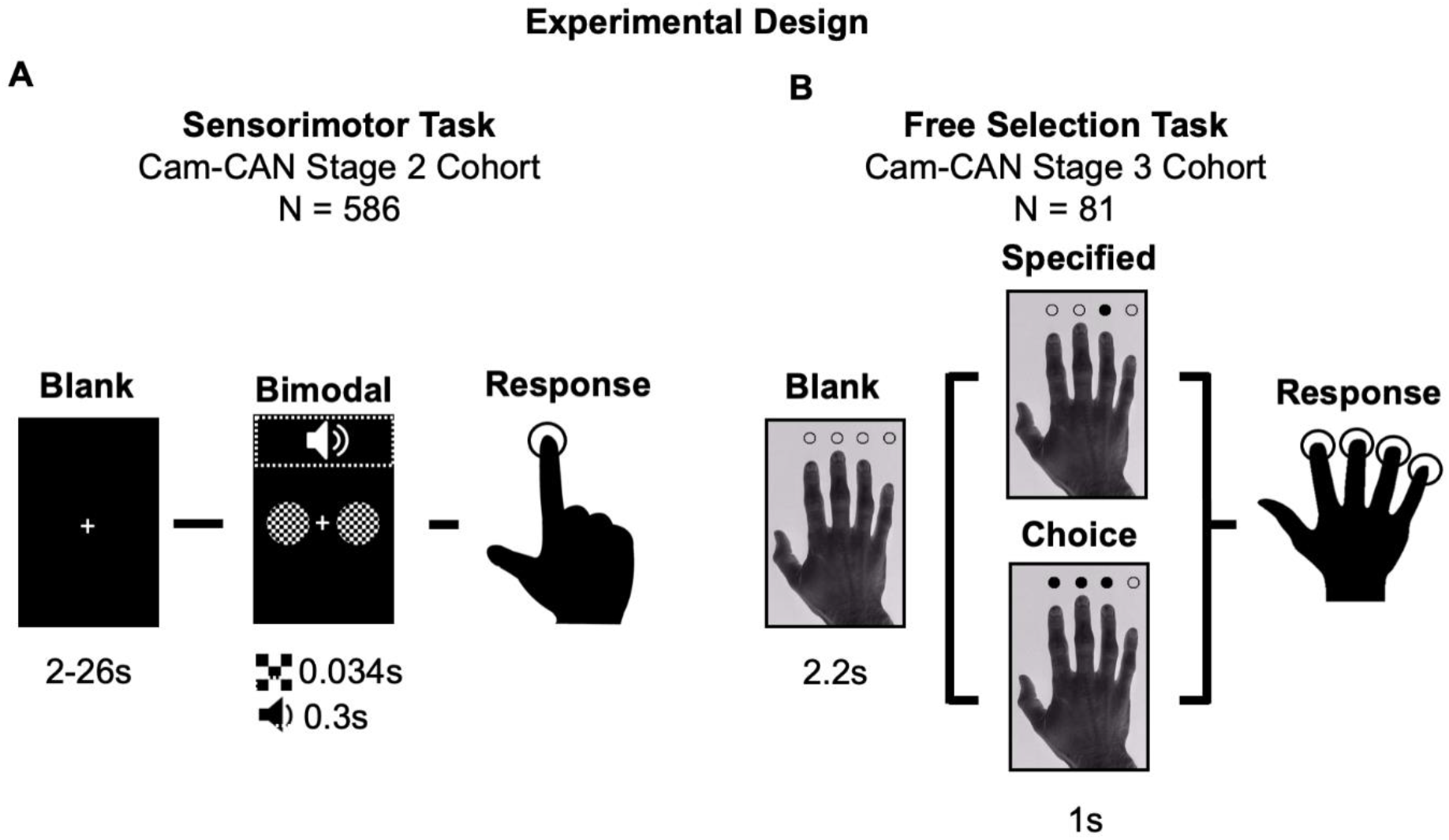
Experimental Design. **(A) Experiment 1**. Sensorimotor task trials began with a blank fixation screen, followed by a bimodal (i.e., audio and visual) stimulus. Participants made finger press responses if they sensed either or both types of stimulus (there were also rare unimodal stimuli on ∼6% of trials, not shown here nor analysed below, in which only an audio or visual stimulus was presented, whose purpose was just to ensure that both modalities needed to be attended). **(B) Experiment 2**. Free selection task trials began with a picture of a hand with circles above the index, middle, ring and little finger. Participants responded with a single finger press that matched one of the cued digits, where only one digit was cued in the “specified” condition, whereas in the “choice” condition, participants were free to choose one of the subset of three digits cued. Both experiments required right-hand responses only.

## Materials & Methods

### Experiment 1: Sensorimotor Task

#### Participants

A healthy, population-derived adult lifespan human sample (N=649; ages approximately uniformly distributed from 18 – 87 years; females = 327; 50.4%) was collected as part of the Cam-CAN study (Stage 2 cohort; Shafto et al., 2014). Participants were fluent English speakers in good physical and mental health based on the Cam-CAN cohort’s exclusion criteria, which excluded volunteers with a low Mini Mental State Examination (MMSE) score (≤ 24), serious current medical or psychiatric problems or poor hearing or vision, as well as being based on standard MRI safety criteria. From this sample, we excluded participants who had missing behavioural measures from either in-scanner (N=4) or out-of-scanner (N=44). We also excluded participants who responded to <90% of trials either in-scanner (N=10) or out-of-scanner (N=5). Thus, the analysed sample consisted of 586 participants (females = 292; 49.8%). The study was approved by the Cambridgeshire 2 (now East of England–Cambridge Central) Research Ethics Committee. Participants gave informed written consent.

#### Materials & Procedure

The sensorimotor task involved 120 bimodal audio/visual trials, as well as eight unimodal trials (four visual and four auditory; Figure 1A) that were included to discourage strategic responding to one modality only. Bimodal trials consisted of visual checkerboards being presented either side of a central fixation (34ms duration) concurrently with a binaural auditory tone (300ms duration). Unimodal trials consisted of either the isolated auditory or visual stimulus. The auditory tones were one of three equiprobable frequencies (300Hz, 600Hz or 1200Hz), which was not relevant to the task or current hypotheses. Participants were instructed to button-press with the right hand index finger when they heard or saw any stimuli. Each trial followed a fixation-only screen with a minimal stimulus onset asynchrony (SOA) of 2 seconds (resulting in SOAs ranging from 2-26 seconds) designed to optimise the estimation of the fMRI impulse response through a sequence of stimulation and null trials (see Shafto et al., 2014).

#### Imaging Data Acquisition & Preprocessing

The MRI data were collected using a Siemens 3T TIM TRIO system with a 32 channel head-coil. A T2*-weighted echoplanar imaging (EPI) sequence was used to collect 261 volumes, each containing 32 axial slices (acquired in descending order) with slice thickness of 3.7mm and an interslice gap of 20% (for whole brain coverage including cerebellum; TR = 1970ms; TE = 30ms; flip angle = 78°; FOV = 192mm x 192mm; voxel-size 3 x 3 x 4.44mm). Higher resolution (1mm x 1mm x 1mm) T1- and T2-weighted structural images were also acquired (to aid registration across participants).

MR data preprocessing and univariate analysis were performed with SPM12 software (Wellcome Department of Imaging Neuroscience, London, www.fil.ion.ucl.ac.uk/spm), release 4537, implemented in the AA 4.0 pipeline (Cusack et al., 2015) described in Taylor et al. (2017). Specifically, structural images were rigid-body registered to an MNI template brain, bias corrected, segmented, and warped to match a grey matter template created from the whole CamCAN Stage 2 sample using DARTEL (Ashburner, 2007; Taylor et al., 2017). This template was subsequently affine transformed to standard Montreal Neurological Institute (MNI) space. The functional images were spatially realigned, interpolated in time to correct for the different slice acquisition times, rigid-body coregistered to the structural image, transformed to MNI space using the warps and affine transforms from the structural image, and resliced to 3mm x 3mm x 3mm voxels.

#### Univariate Imaging Analysis

To estimate activity for univariate voxelwise contrasts (i.e., to define ROIs), five conditions (i.e., 3 bimodal conditions, one per tone frequency, and 2 catch conditions, per audio or visual format) were distinguished within a general linear model (GLM) for each participant using SPM. A regressor for each condition was created from delta functions, aligned to the onset of a stimulus, that were convolved with SPM’s canonical hemodynamic response function, plus its temporal and dispersion derivatives, resulting in three regressors per condition. The null events were excluded from the model, and therefore, all regression coefficient were defined relative to this baseline activity. Six additional regressors representing the three rigid body translations and rotations estimated in the realignment stage were included in each GLM to capture residual movement-related artifacts. Finally, the data were scaled to a grand mean of 100 over all voxels and scans within a session, and the model was fit to the data in each voxel. The autocorrelation of the error was estimated using an AR(1)-plus-white-noise model, together with a set of cosines that functioned to high-pass filter the model and data to 1/128 Hz, that were estimated using restricted maximum likelihood. The estimated error autocorrelation was then used to “prewhiten” the model and data, and ordinary least squares used to estimate the model parameters. Contrasts were used to average across the 3 tone frequencies in the bimodal trials (i.e., the rarer unimodal trials were not analysed further). This model was used for ROI definition and MVB, whereas for regressions involving univariate data, we used a least squares separate (LSS) approach (Abdulrahman & Henson, 2016) before averaging over voxels.

#### Behavioural Measures

Reaction time (RT) was the time from stimulus onset to button press onset. RTs were estimated during the fMRI sensorimotor task (i.e., in-scanner RT) and during an independent lab-based simple RT task (i.e., out-of-scanner RT) performed during Stage 1 of the Cam-CAN project. In the out-of-scanner task, participants were presented with the same picture stimulus as the free selection experiment (Figure 1B; see section: Materials & Methods - Experiment 2: Free Selection - Materials & Procedure) where, for each trial (N = 50), a blank circle above an index finger was filled black, cueing a button-press response to be performed as quickly as possible. Upon pressing the button (or after 3s), the circle’s fill was cleared and followed by pseudo-random inter-trial interval (see Shafto et al., 2014). Note that, while the out-of-scanner task was speeded, the in-scanner task was unspeeded (so that older participants did not feel too challenged). For each participant, both the mean and standard deviation (variability) of RTs across trials were computed.

### Experiment 2: Free Selection

#### Participants

Participants were a subset of the cohort in Experiment 1 who also completed the Free Selection fMRI experiment during Stage 3 of Cam-CAN data collection (N=87; approximately uniformly distributed from 19 - 85yrs; females = 38; 43.7%). We excluded 6 participants whose out-of-scanner RT measures were not collected (all remaining participants responded to >90% of trials, and were correct for >75% of trials). Therefore, the analysed sample consisted of 81 participants (females = 35).

#### Materials & Procedure

The free selection task was adapted from the 3-choice free selection task of Zhang et al. (2012), which involves a visually-paced right hand button press task that is typically used to examine executive control and action decisions in ageing. Across 240 trials, participants were presented with an image of a right hand and pressed a button with one finger in response to a cue (see Figure 1B). Individual trials involved either one (“specified” condition; N=120, split equally between each of the four fingers) or three (“choice” condition; remaining 120 trials) of the circles being filled black. In both cases, participants were instructed to respond as quickly as possible with a single button-press from a cued digit; thus for choice trials the responding finger could be freely selected. Cues were pseudorandomly ordered so that participants do not see four or more responses of the same condition in a row (see Shafto et al., 2014 for more details). A short gap (either 4.2s or 6.2s) separated blocks of 20 trials.

#### Imaging Data Acquisition & Preprocessing

Data acquisition and processing were the same as in Experiment 1 (see section: Methods - Experiment1: Sensorimotor Task - Imaging Data Acquisition & Preprocessing), aside from an increased number of volumes being acquired (296) due to a longer session duration.

#### Univariate Imaging Analysis

The procedure described for Experiment 1 was repeated here (see section: Materials & Methods - Experiment 1: Sensorimotor Task - Univariate Imaging Analysis), except that only the canonical HRF was used (because the blocked nature of trials prevents reliable estimation of the HRF derivatives; Henson, 2015). For the present analyses, we combined onsets across the specified and choice conditions, leaving four predictors based on which finger was pressed (i.e., index, middle, ring and little). These four conditions were averaged to estimate the mean response versus baseline.

#### Behavioural Measures

The same variable definitions and computations were used as described for Experiment 1 (see section: Materials & Methods - Experiment 1 - Sensorimotor Task). Unlike Experiment 1, the out-of-scanner RT variables were measured during a choice RT task that had a design more comparable to the in-scanner free selection task. Specifically, the choice RT task had the same parameters as the simple RT task, but on each trial (N=67) any one of the four circles above the fingers could be filled black, and the participant was instructed to press the corresponding finger as quickly as possible.

## General Methods

### Regions of Interest (ROIs)

A standard group univariate voxelwise approach was used to define a contralateral sensorimotor cortex ROI, based on contrasting all bimodal trials versus baseline in Experiment 1. Specifically, the 70 most significant voxels (based on t-statistic rank) were selected according to the peak closest to the left “hand knob” landmark in the central sulcus (Yousry et al., 1997; Figure 2A; for MNI coordinates see Figure 2A; Table 1). This contralateral ROI was mirror flipped (i.e., x-coordinate reversed in sign) to create the ipsilateral sensorimotor cortex ROI (Figure 2A; Table 1). Note that this ROI selection based on the average response versus baseline is averaged across age (i.e., not biased to show age effects). The same ROIs were applied to Experiment 2 for consistency. Note that images were spatially smoothed (10mm Gaussian kernel) for the purpose of ROI definition only. All ROI analyses used unsmoothed data.

**Figure 2.**
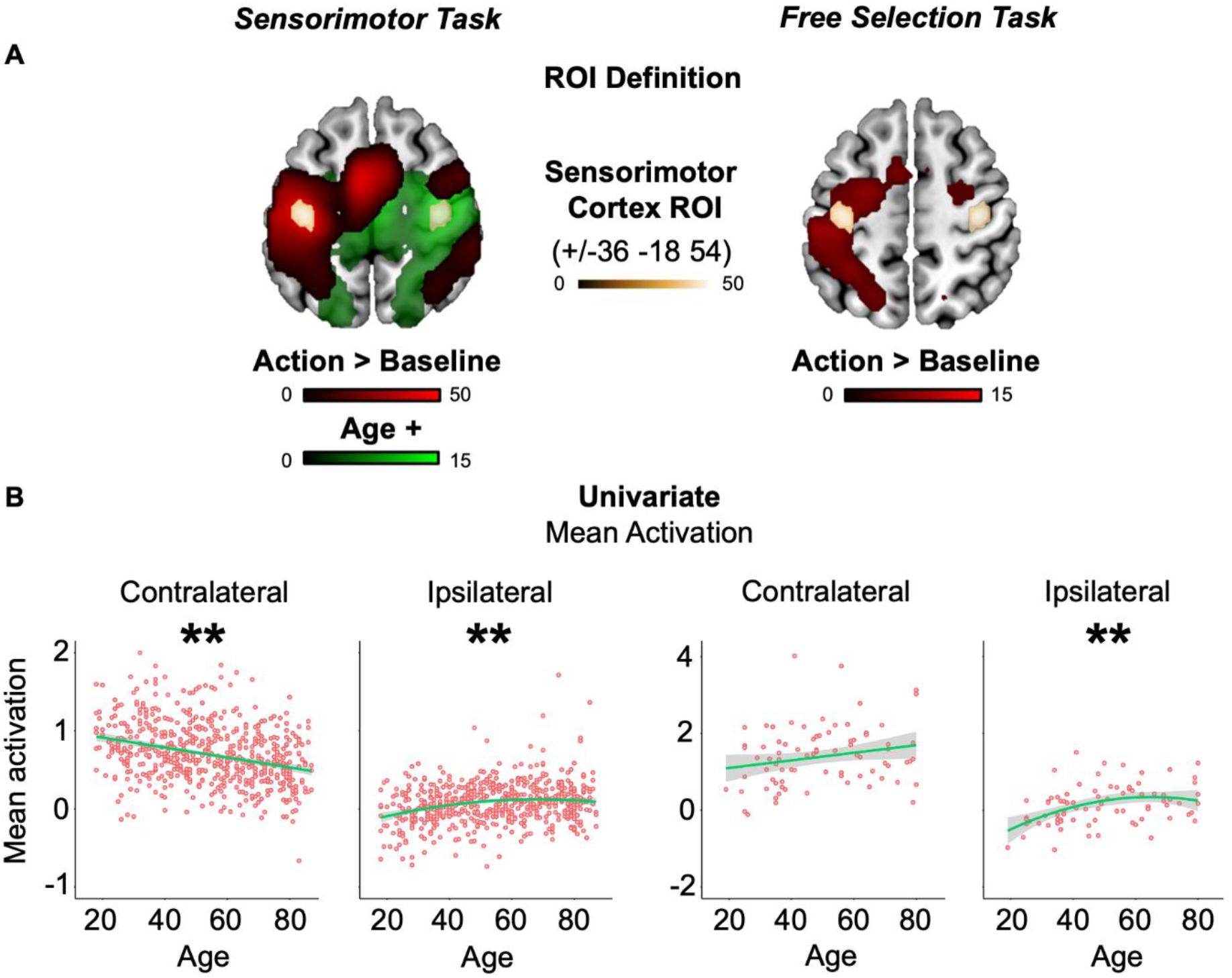
ROI definitions and responses. **(A) ROI Definitions**. Univariate whole-brain voxelwise T-tests are displayed on a standard template brain for all actions > baseline (red map) and for positive (linear) effect of age (green map). Color depth indicates T-statistic value. The actions > baseline contrast from Experiment 1 was used to define the functional ROI in sensorimotor cortex (gold map), which was mirror-flipped across hemispheres for unbiased analysis of age effects in both experiments (see section: Methods - General Methods - ROIs). **(B) Univariate ROI Responses**. Consistent with HAROLD, increased age predicted increased univariate activation of the ipsilateral ROI in both experiments, accompanied by the opposite pattern in the contralateral ROI. Green lines represent robust-fitted regression lines (with a second polynomial expansion in cases where a significant quadratic component was observed) and shaded areas show 95% confidence intervals. [* = p < 0.05, ** = p < 0.01].

**Table 1.** Age effects on mean univariate and spread of multivariate action effects.

| Experiment | Measure/ROI | Age Effect |  | Linear |  | Quadratic |  |
| --- | --- | --- | --- | --- | --- | --- | --- |
|  |  | F | p | t | p | t | p |
| Sensorimotor |  |  |  |  |  |  |  |
|  | Univariate Mean |  |  |  |  |  |  |
|  | Contralateral | 31.7 | <.0001 | -7.95 | <.0001 | -0.3 | 0.76 |
|  | Ipsilateral | 18.2 | <.0001 | 5.26 | <.0001 | -3.11 | .002 |
|  | Multivariate Spread |  |  |  |  |  |  |
|  | Contralateral | 4.82 | .008 | 2.31 | .021 | -2.08 | .038 |
|  | Ipsilateral | 2.97 | .052 | - | - | - | - |
| Free Selection |  |  |  |  |  |  |  |
|  | Univariate Mean |  |  |  |  |  |  |
|  | Contralateral | 4.49 | .014 | 1.88 | .067 | -2.32 | .025 |
|  | Ipsilateral | 9.31 | .0002 | 3.66 | .0004 | 0.900 | .032 |
|  | Multivariate Spread |  |  |  |  |  |  |
|  | Contralateral | 2.83 | .065 | - | - | - | - |
|  | Ipsilateral | 2.02 | .14 | - | - | - | - |

### Multivariate Bayesian Decoding (MVB)

A series of MVB decoding models were fit to assess the information about actions represented in each ROI or combination of ROIs. Each MVB decoding model is based on the same design matrix of experimental variables used in the univariate GLM, but the mapping is reversed: many physiological data features (derived from fMRI activity in multiple voxels) are used to predict a psychological target variable (Friston et al., 2008). This target (outcome) variable is specified as a contrast. In both experiments, the outcome was whether an action had been performed (versus baseline), with all covariates apart from those involved in the target contrast (i.e., the null space of the target contrast) removed from both target and predictor variables.

Each MVB model was fit using a parametric empirical Bayes approach, in which empirical priors on the data features (voxelwise activity) are specified in terms of spatial patterns over voxel features and the variances of the pattern weights. As in earlier work, we used a “sparse” spatial prior in which patterns are individual voxels. Because these decoding models are normally ill-posed (with more voxels relative to scans, or more precisely, relative to degrees of freedom in the timeseries), these spatial priors on the patterns of voxel weights regularise the solution. MVB also uses an overall sparsity (hyper) prior in pattern space that embodies the expectation that a few patterns make a substantial contribution to the decoding and most make a small contribution.

The pattern weights specifying the mapping of data features to the target variable are optimized with a greedy search algorithm using a standard variational scheme, which iterates until the optimum set size is reached (Friston et al., 2007). This is done by maximizing the free energy, which provides an upper bound on the Bayesian log evidence (the marginal probability of the data given that model). The evidence for different models predicting the same psychological variable can then be compared by computing the difference in their log evidences (equivalent to the log of the “Bayes Factor”; Friston et al., 2008; Chadwick et al., 2012; Morcom & Friston, 2012). In this work, the main outcome measures were the log evidence for each model and the spread (standard deviation) of weights across voxels in the ROI (Morcom & Henson, 2018).

To test whether ipsilateral activity was compensatory, we used a “boost” measure (Morcom & Henson, 2018) to assess the contribution of the ipsilateral ROI to performing actions. This used Bayesian model comparison within participants to assess whether a combined contralateral-ipsilateral (i.e., bilateral) model boosted prediction of actions relative to a contralateral-only model. The compensatory hypothesis, in which the ipsilateral hemisphere is engaged to a greater degree in older age and improves performance, predicts that a boost will be more often observed with increasing age. The dependent measure was the log model evidence coded categorically for each participant to indicate the outcome of the model comparison. The three possible outcomes were as follows: a boost to model evidence for bilateral relative to contralateral-only models (difference in log evidence > 3), ambiguous evidence for the two models (−3 < difference in log evidence < 3), or a reduction in prediction of action for bilateral relative to contralateral-only (difference in log evidence < -3). These values were chosen because a log difference of 3 corresponds to a Bayes Factor >20 which is generally considered strong evidence (Lee & Wagenmakers, 2014).

For the across-participant analyses of this MVB “boost”, participants were only included if their data allowed reliable decoding by the bilateral model (Morcom & Henson, 2018). To determine this, we contrasted the evidence for the bilateral model with that from models in which the design matrix (and therefore the target variable) was randomly phase-shuffled. One-tailed t-tests were used to compare whether the mean difference between true and shuffled differences in log-evidence was greater than 3 (Morcom & Henson, 2018; Figure 4A), which left a total of 650 and 54 participants for Experiment 1 and 2, respectively (i.e., N=4 and N=27 excluded respectively). For additional control analyses, we repeated the MVB boost analysis with models where voxel sizes were equated (see Results section). For one of the control analyses which involved halving the number of voxels in the bilateral model, we repeated this preliminary phase shuffling step (because a different bilateral model was used) which led to excluding four additional participants in Experiment 1 and prevented this particular control analysis for Experiment 2, because there was not evidence that decoding was possible from this ROI (*p* = 0.12).

### Experimental Design & Statistical Analysis

Age effects on continuous univariate, behavioural and multivariate measures were tested using robust regression in R [version: 3.6.1] with the *rlm* function (package: MASS [version: 7.3-51.4]), in order to down-weight extreme values (Venables & Ripley, 2002). These regression analyses used standardised linear and quadratic age predictors. Two-tailed robust F-tests (Wald tests) were used to test the significance of regression coefficients: we first tested for general age effects (linear and/or quadratic), and if significant (α level of 0.05), we performed post-hoc Wald tests on linear and quadratic age predictors separately. Analysis of the categorical outcomes for the between-region MVB model comparison (Figure 4B) used ordinal regression. When all three categorical outcomes were observed, this was implemented with the *polr* function (package: MASS; as in Henson & Morcom, 2018; Table 4), whereas *glm* (package: stats [version: 3.61]) was used in binary cases (i.e., when ‘reduction’ was not observed for any participant; Figure 4B). For ordinal regression, the results are reported from a model containing only the linear age term, due to the categorical nature of the data (though the same pattern of findings were observed with the full quadratic model; see Table 4 with chi-square tests for general age effects).

**Table 2.**
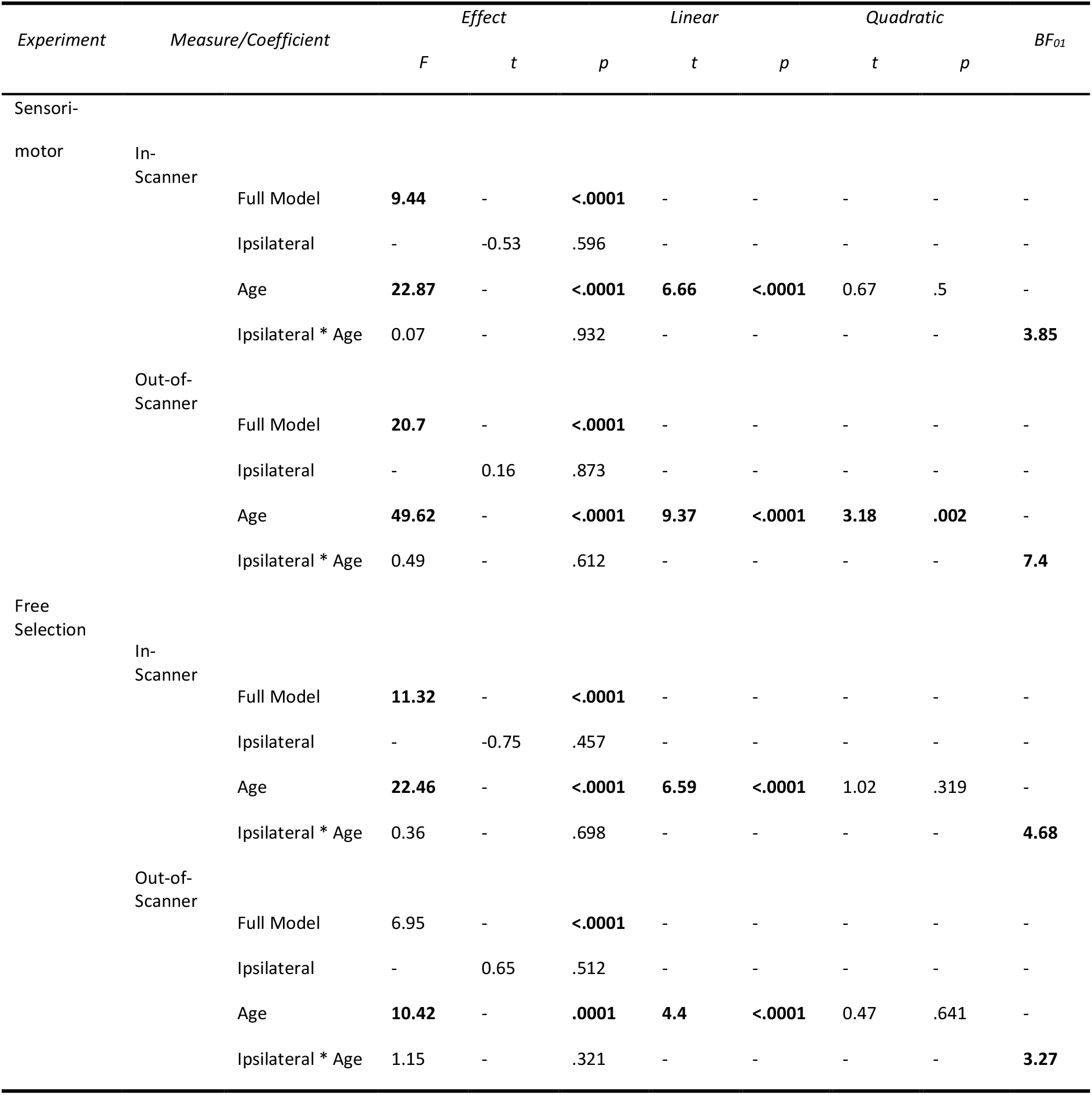
Age and mean univariate effects from behavioural multiple regression with RT variability. Degrees of freedom for Experiment 1: full model *F*(5,580), age effect *F*(2,580) and *t*(580); Experiment 2: full model *F*(5,75), age effect *F*(2,75) and *t*(75). Bold text indicates *p* < 0.05.

**Table 3.**
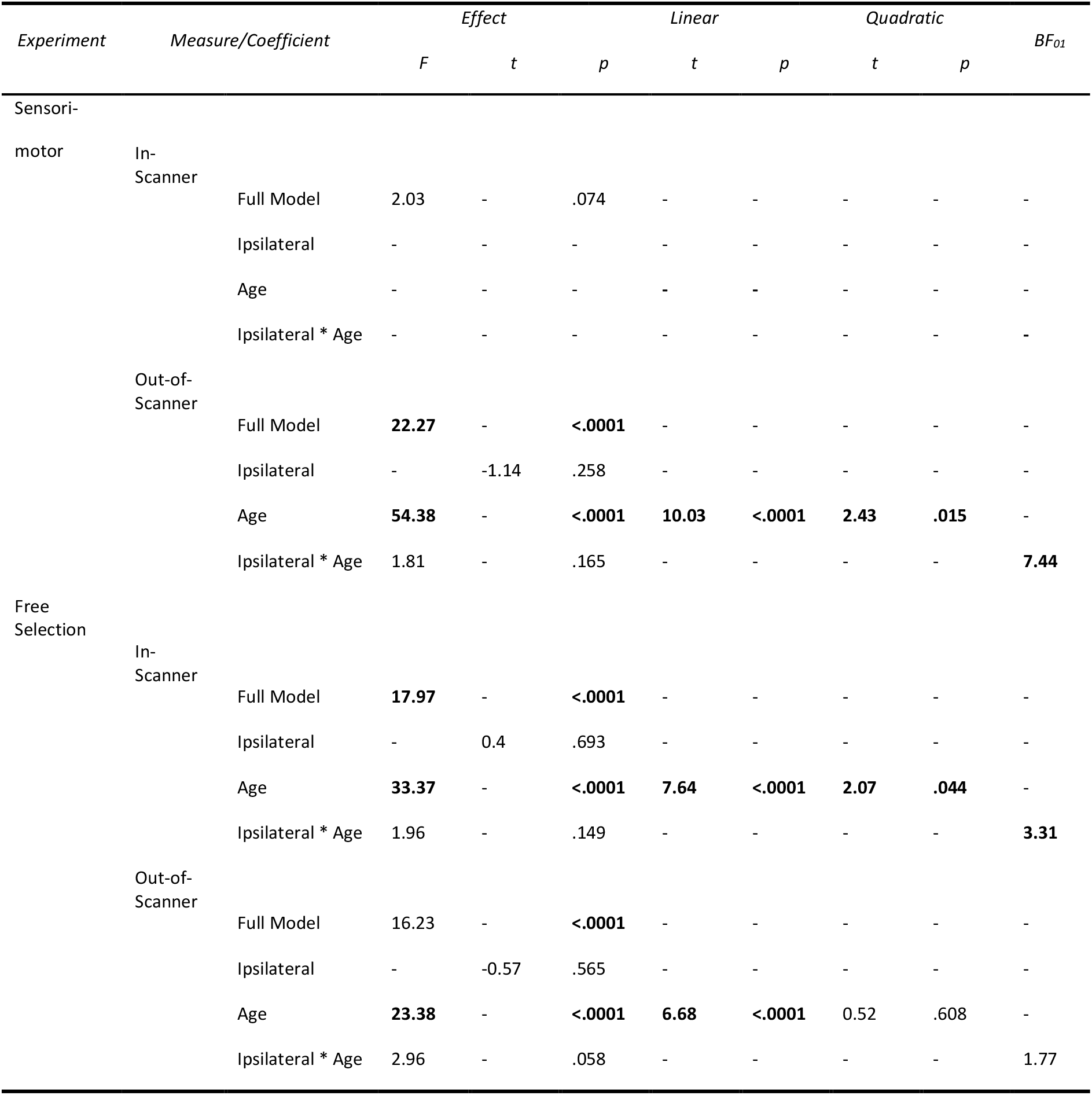
Age and mean univariate effects from behavioural multiple regression with mean RT. See Table 2 for degrees of freedom and conventions.

**Table 4.**
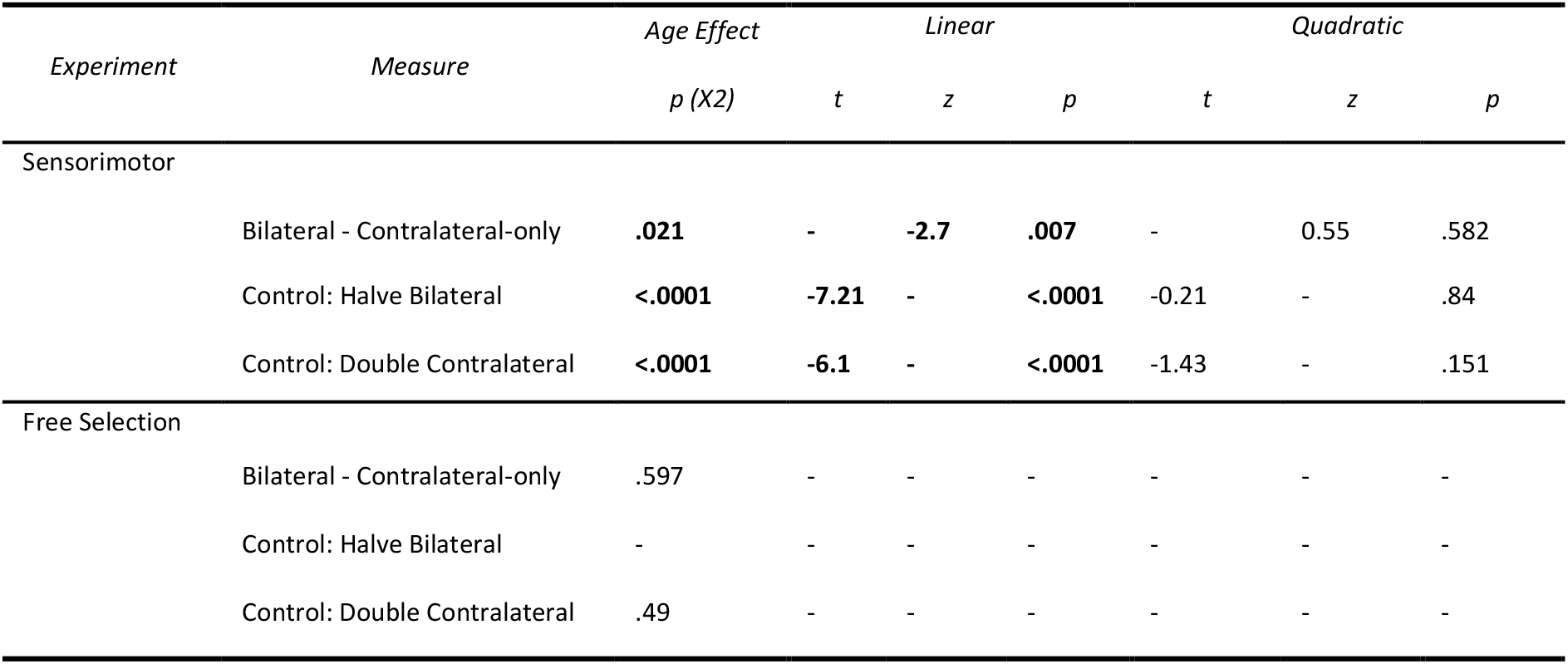
Age effects from ordinal regression during the MVB boost analyses.

When important, null-hypothesis significance tests were supplemented with Bayes factors (Wagenmakers, 2007; Rouder et al., 2009). For continuous outcomes, we used the *lmbf* (package: BayesFactor [version: 0.9.12-4.2]) with default parameters (Rouder, et al., 2012) to contrast models with/out the effect predicted by compensation accounts. For categorical outcomes (i.e., MVB model comparison), we used the *brm* function (package: brms [version: 2.10.0]) with the Bernoulli family function to test for the absence of the hypothesis predicted by compensation (i.e., age effect > 0). A Student’s t distribution prior was used, based on 7 degrees of freedom, a mean of 0, and a scale factor of 10 and 1 for the intercept and slope, respectively (e.g., Wagenmakers et al., 2010). The Bayes Factors were interpreted according to criteria set out by Jeffreys (1961; cited from Jarosz & Wiley, 2014), where a BF_01_ between 1-3, 3-10 and >10 indicates ‘anecdotal’, ‘substantial’ and ‘strong’ evidence in favour of the null, respectively.

### Data Availability

Raw and minimally pre-processed MRI (i.e., from automatic analysis; Taylor et al., 2017) and behavioural data are available by submitting a data request to Cam-CAN (https://camcan-archive.mrc-cbu.cam.ac.uk/dataaccess/). The univariate and multivariate ROI data, and behavioural data, can be downloaded from the Open Science Framework alongside analysis code (https://osf.io/seuz5/).

## Results

### HAROLD univariate effect

The univariate voxel-wise contrast of key press versus baseline during the sensorimotor task (Experiment 1), averaged across participants, showed strong contralateral activation throughout fronto-parietal cortex (red voxels in Figure 2A, left). The x-coordinates of a contralateral motor cortex ROI that spanned suprathreshold voxels in the pre-central gyrus were flipped to define an ipsilateral motor cortex ROI (Figure 2A gold voxels; Table 1; see Methods - General Methods - ROIs).

Consistent with HAROLD’s predictions, when averaging over voxels within the ipsilateral ROI, there was a significant effect of age on univariate activity, with an increase in activation that flattened off in old age (Figure 2B, left), in line with significant linear and quadratic components (see Table 1). In fact, although the ROI was defined independently of age, it entirely overlapped voxelwise results from a positive t-contrast on the (linear) effect of age (green voxels in Figure 2A, left). The significant age effect for the contralateral ROI was in the opposite direction, with mean activity decreasing linearly as a function of age (Figure 2B, left; Table 1).

When applying these ROIs defined in Experiment 1 to Experiment 2, we replicated this HAROLD effect, where greater age was associated with greater ipsilateral sensorimotor cortex activation; an age effect that again decelerated in later life (Figure 2B, right; Table 1). Unlike Experiment 1, no suprathreshold age effect was observed when repeating the voxelwise linear contrast, possibly owing to the lower statistical power than in Experiment 1 (Figure 2A, right). Again, the trend for the age effect in the contralateral ROI was in the opposite direction, though only the quadratic term was significant when tested independently (Table 1).

### Testing Compensation: Behavioural

If the increased univariate activity in ipsilateral sensorimotor cortex is compensatory, it might be expected to benefit task performance. We measured task performance using the variability (Table 2) and mean (Table 3) of RTs for both the in-scanner and out-of-scanner motor tasks. First, we tested whether there was a main effect of age on RT (Figure 3A). For variability of simple RTs in Experiment 1, significant effects were found for RTs recorded both in-scanner and out-of-scanner, where higher ages were linearly associated with increased variability, that is, worse performance (Table 2; Figure 3A left). These significant age effects were replicated in the choice RTs of Experiment 2, both in-scanner and out-of-scanner, where, again, a linear positive change in RT variability was predicted by increased age (Table 2; Figure 3A right). For the out-of-scanner measure in Experiment 1, the quadratic component was also significant, such that the increase in RT variability actually accelerated in old age (Figure 3A). For mean simple RTs, there was no significant effect of age for the in-scanner measure during Experiment 1 (Table 3; Figure 3A left), most likely because this version of the task was not speeded. However, there was an age effect on the speeded out-of-scanner task, with, like for RT variability, significant linear and quadratic components, indicating that worse performance accelerated in old age (Table 3; Figure 3A left). For Experiment 2, the results for mean RT were similar to those reported for RT variability (i.e., there was a positive linear effect of age; Figure 3A right).

**Figure 3.**
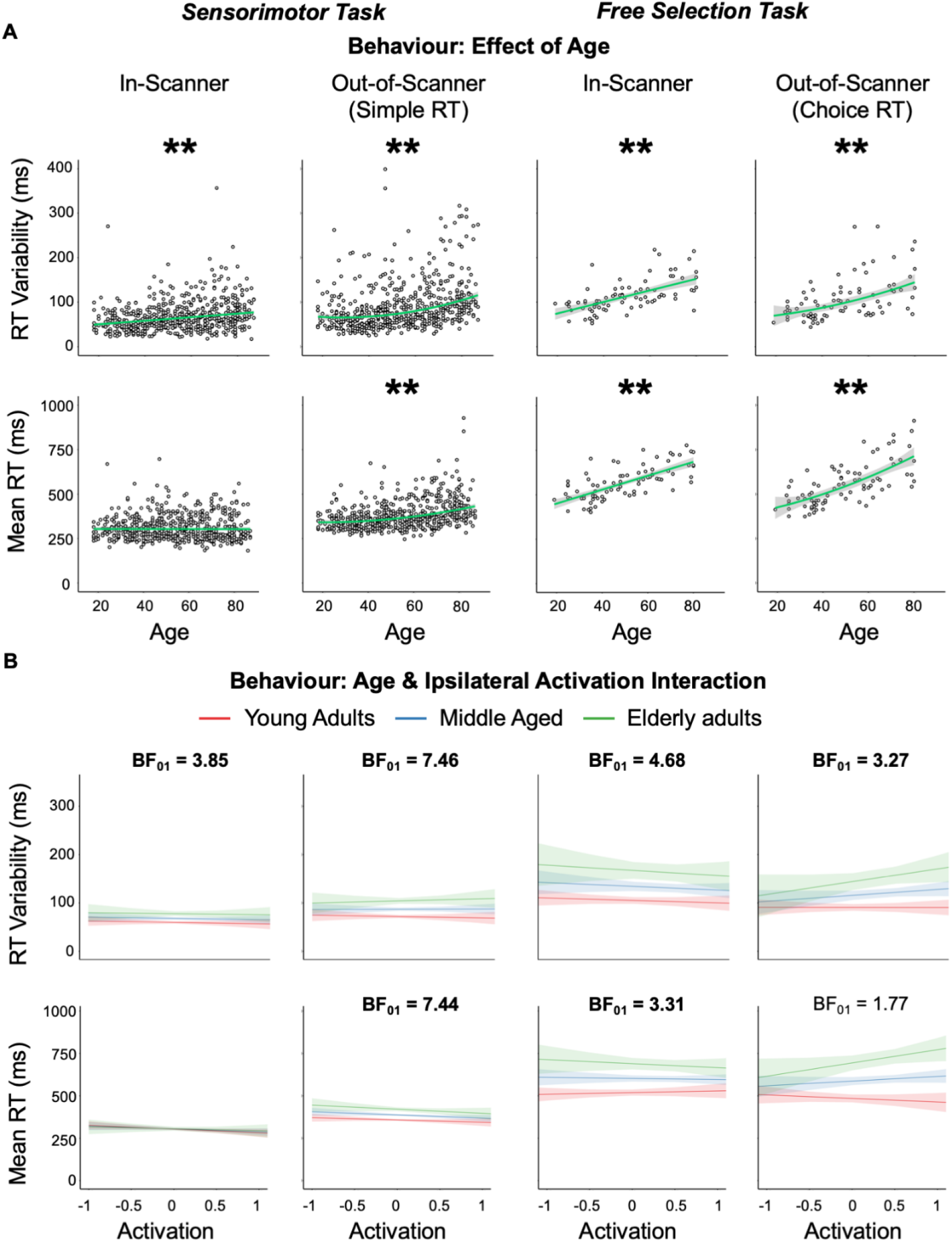
Behavioural results for RT variability and mean RT. **(A) Effect of age**. Increased age predicted worse performance (greater RT variability/mean) across Experiments 1 (left) and 2 (right) whether measures were acquired in- or out-of-scanner. For asterisk and regression line conventions, see Figure 2. **(B) Interaction between age and ipsilateral univariate activation**. No significant interactions between age and ipsilateral mean activation were observed across experiments, regardless of whether measures were acquired in-or out-of-scanner. Bayes Factors for the null (BF_01_) that had substantial evidence for this lack of interaction are in bold. Though the interaction was tested in a continuous fashion, tertile splits were used to define age groups (red, blue, green lines) for purposes of illustration.

Having established age effects on task performance, the critical question was whether this age-related variance was related to ipsilateral motor activation, with compensation predicting that higher activation in older people would relate to better (i.e., faster and less variable) RTs. To assess this, we used multiple regression to test whether age, ipsilateral activation and their interaction predicted RT variabilty. If ipsilateral activity is compensatory and has an overall benefit to performance, then one would expect a significant interaction between age and ipsilateral activity, whereby the tendency for higher ipsilateral activation to be associated with reduced RT variability would increase with age. However, contrary to this prediction, no significant interaction between ipsilateral activity and age was observed when predicting RT variability (Table 2; Figure 3B top row) or mean RT (Table 3; Figure 3B bottom row) either in- or out-of-scanner for Experiment 1 or Experiment 2. In fact, Bayes Factors presented consistent evidence in favour of no interaction for all measures with a significant age effect in Experiment 1 (Figure 3B left) and three of the four measures in Experiment 2 (Figure 3B right) (see Table 2, 3).

Lastly, we tested the possibility that ipsilateral recruitment in later life partially compensates for reduced contralateral function. Although compensatory recruitment may have a net benefit to performance, compensation can also function like a walking stick, being engaged to a greater degree by people with a greater need for it (Backman, 1985). In such cases, compensatory brain activity may correlate negatively with individual performance in older adults (i.e., only partially, rather than fully, compensating relative to younger people (e.g., Daselaar & Cabeza, 2005; de Chastelaine et al., 2011; Morcom & Johnson, 2015)). We therefore used multiple regression to ask whether ipsilateral activation would relate positively to performance once effects of contralateral impairment were taken into account by including contralteral mean activity (i.e., degree of impairment) as a predictor. We expected that ipislateral activation would be associated with better performance only in people with low contralateral activity, not in people with maintained (high) contralateral activity, who did not need to compensate. This type of compensatory account therefore predicts an interaction between contralateral and ipsilateral activity in relation to RT performance. To test this, we repeated analyses replacing the age predictor with contralateral mean activity. In neither experiment was there a signficant interaction between ipsilateral activity and contralateral activity (all *p’s* ≥ 0.074). This remained the case even if we added age as a third predictor. Indeed, there was substantial Bayesian evidence against this effect for all measures (with or without age) in Experiment 1 (BF_01_’s ≥ 3.19) and, in Experiment 2, for the in-scanner RT variability measure (3 predictor model: BF_01_ = 4.04; all other BF_01_*’s* ≤ 2.98).

### Testing Compensation: Multivariate

We further tested the compensation account of HAROLD using a multivariate approach. If the increasing ipsilateral activation with age reflected compensation, then multivoxel analyses should show that this increased activity carries additional information about actions, over and above that provided by the contralateral hemisphere. Note that this could happen even if the mean response across voxels did not relate to behavioural performance, as in the previous section (e.g., Morcom & Henson, 2018).

To test this, we first applied MVB to the combination of contra- and ipsilateral motor ROIs (i.e., 138 voxels in total), to check classification of an action was above chance, by comparing real versus phase-shuffled fMRI data. Results showed that the difference in log model evidence was greater than 3 on average across participants in both Experiment 1 (*t*(585) = 44.27, *p* < 0.0001) and Experiment 2 (*t*(80) = 4.57, *p* < 0.0001). Figure 4A shows that decoding was possible for the majority of participants. There was also a significant linear effect of age on the probability that model evidence was (or was not) greater than 3 for Experiment 2 where successful decoding was more likely to occur for older ages (*z*(80) = 3.11, *p* = 0.005). In Experiment 1, this was not examined due to the rarity (N=4) that the difference in model evidence was less than 3 (Figure 4A).

**Figure 4.**
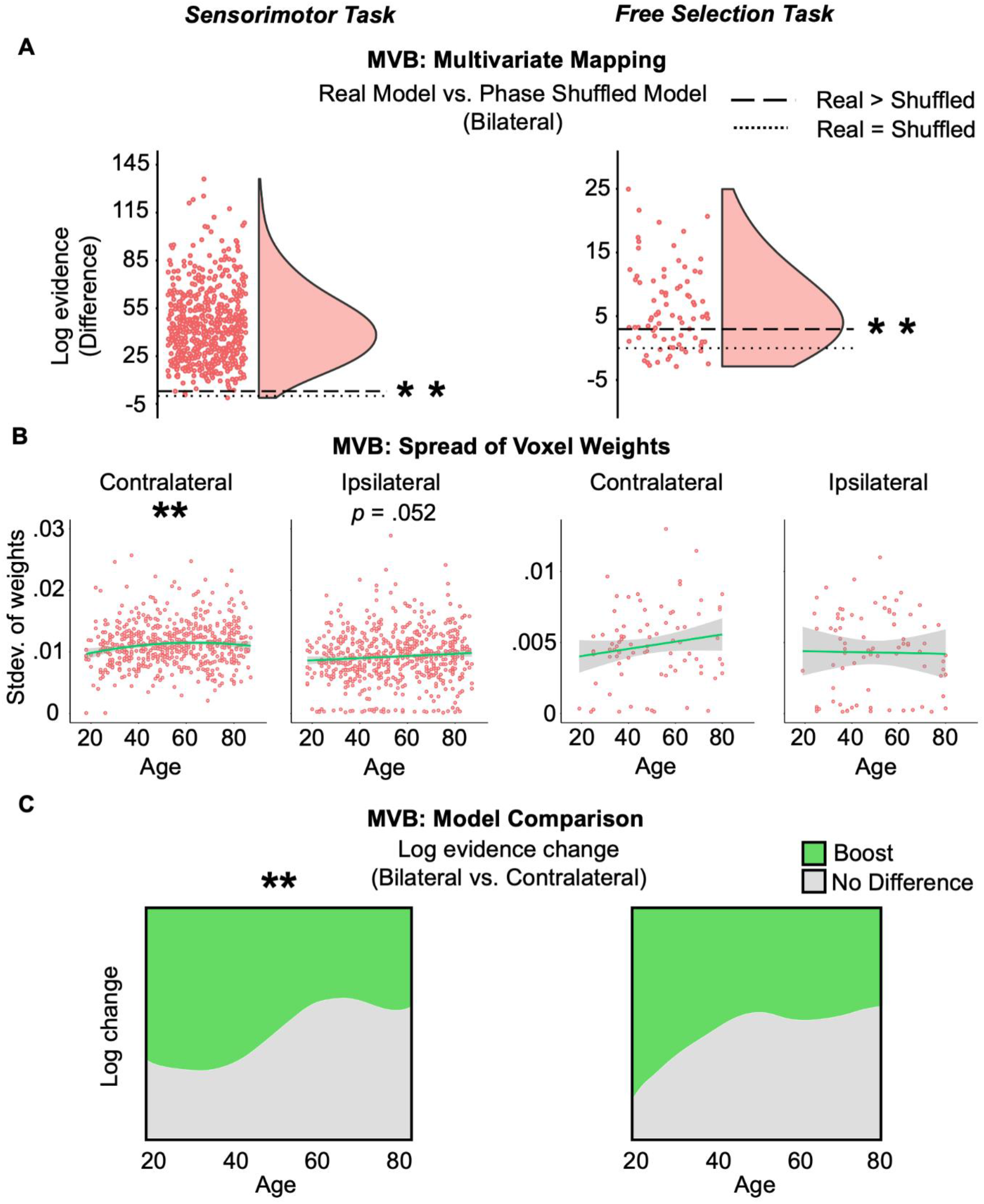
MVB results. **(A) Multivariate Mapping**. For the target outcome being decoded (i.e., performing an action), the difference in log model evidence was significantly higher than 3 (dashed line) when using real (as opposed to phase-shuffled) action onsets, indicating reliable decoding across both experiments (left and right; dotted line indicates a difference of 0). **(B) Multivariate ROI Responses**. The spread of voxel weights showed an increase with age in the contralateral ROI in Experiment 1, plus a similar trend in Experiment 2, and for the ipsilateral ROI in Experiment 1, but not in Experiment 2. **(C) Model Comparison**. For Experiment 1 (left) results showed that, contrary to a compensatory account, increased age actually led to a reduction in the likelihood of a boost when including ipsilateral voxels. For the free selection task (right), the effect of age was in the same direction but did not reach significance.

Having shown that MVB decoding was possible, one measure of multivariate information is the spread (e.g., standard deviation) of voxel classification weights (Morcom & Henson, 2018). This measure indexes the absolute magnitude of unique voxel contributions to the task. We therefore calculated spread for MVB models applied to each ROI separately. The results are shown in Figure 4B. In Experiment 2, no significant effect of age was observed on the spread of either the contralateral or ipsilateral weights (Figure 4B right; Table 1). However, in Experiment 1, there was a significant effect of age on spread for the contralateral ROI, in which the linear and quadratic components were significant, indicating that decodable information about a right finger press increased with age (in a decelerating fashion) in contralateral sensorimotor cortex (Figure 4B left; Table 1). The effect of age on weight spread was not significant for the ipsilateral hemisphere, though there was a trend in the same direction (Figure 4B left; Table 1). Thus, unlike in Morcom & Henson (2018), it might be that multivariate information about a right finger press increases with age in ipsilateral motor cortex, the region that is proposed to compensate. However, even if this age-related increase occurs for both ipsilateral and contralateral ROIs, it is possible that the same information is being represented in each hemisphere. That is, any age-related increase in information in the ipsilateral ROI could be redundant with that in the contralateral ROI, rather than being unique (i.e., compensatory).

Therefore the crucial test was whether the information in the ipsilateral ROI improved action prediction compared to that in the contralateral ROI. Using MVB in Experiment 1, the proportion of participants showing such an ipsilateral “boost” actually decreased, rather than increased, with age (linear, *z*(580) = -2.86, *p* = 0.004) (Figure 4C). In other words, contrary to a compensatory account, the odds that model evidence was boosted by including ipsilateral with contralateral activity for older adults was 0.61 times that for younger adults. Indeed, the Bayes Factor provided strong evidence in favour of accepting the null over the compensatory hypothesis (BF_01_ = 21.99). For Experiment 2, no significant effect of age was found (*z*(52) = -0.88, *p* = 0.38) (Figure 4C) though, in line with Experiment 1, there was substantial Bayesian evidence against the compensatory hypothesis (BF_01_ = 4.87).

We performed a final check where we explicitly matched the number of voxels in the combined versus contralateral models. Regardless of whether we halved the number of voxels in the combined model (from 140 to 70), or doubled the number of voxels in the contralateral model (from 70 to 140), the significant linear negative effect of age in Experiment 1 and non-significant effect in Experiment 2 were replicated (after halving: Experiment 1: *t*(575) = -10.02, *p* < 0.0001; after doubling: Experiment 1: *t*(579) = -14.13, *p* < 0.0001; Experiment 2: *p* = 0.29). All findings were of the same pattern across experiments when models contained both the linear and quadratic age terms (see Table 4).

## Discussion

After replicating univariate HAROLD effects in motor cortex (e.g., Naccarato et al., 2006; Tsvetanov et al., 2015) across two finger movement fMRI experiments in a large lifespan sample, we tested if the additional ipsilateral activation in older adults reflected a compensatory mechanism. None of the behavioural or multivariate measures, in either experiment, showed age effects that would be predicted by a compensation account of HAROLD. In fact, Bayes Factors demonstrated substantial evidence against compensatory interactions between age and ipsilateral mean activation for all behavioural analyses in Experiment 1 as well as for many in Experiment 2. Likewise, the Bayes Factors for the MVB boost analysis were strongly against any age effect, where a compensation account would predict an age-related boost for action decoding with additional ipsilateral voxels. In fact, for Experiment 1, an age effect was observed in the opposite direction: as age increased, adding the additionally activated voxels was found to be *less* likely to improve action decoding.

Previous tests of age-related compensatory accounts have been inconclusive (Ward, 2006). Some of this uncertainty in the literature might owe to differences in sample size, task and/or analysis. At least for finger presses, we believe that our sensorimotor results are more conclusive because: (1) they come from a relatively large and more population-representative samples, (2) they simultaneously model age, behaviour and (ispi- and contra-lateral) activation, and (3) include a Multivariate Bayesian approach that tests not only whether there is multivoxel information about actions in ipsilateral cortex, but also whether this information is distinct (i.e., non-redundant) from that in contralateral cortex.

Another reason for the lack of agreement in the literature is that compensation may take more than one form, and is difficult to test with small samples (for reviews, see Scheller et al., 2014; Morcom & Johnson, 2015). Compensation may not always give rise to a positive relationship between the compensatory activation and behaviour. Instead, it might only be partially successful, analogous to a walking stick that helps older people walk faster than without it, but still not as fast as they would in the absence of any age-related decline (Daselaar & Cabeza, 2005; de Chastelaine et al., 2011). In the present case, if performance declines with age due to reduced contralateral motor function, this may be only partially offset by compensatory ipsilateral activation, giving rise to a net negative association between ipsilateral activity and performance in older people. We therefore tested for this ‘partial compensation’ in additional behavioural analyses that used contralateral activity as a surrogate for the degree of age-related motor impairment. However, there was evidence against the predicted interaction of contralateral and ipsilateral activity on performance. Moreover, partial compensation is inconsistent with our MVB results, where multivariate information was more likely to be unchanged or reduced with the purported compensatory mechanism (i.e., ipsilateral activity) with increasing age.

Thus our multivariate (MVB) analyses arguably provide the strongest evidence against compensation (Figure 4C). This result is consistent with the only other multivariate experiment, to our knowledge, to examine this in the motor system, where MVPA demonstrated less distinctive ipsilateral motor cortex activity with age (Carp et al., 2011). However, our results strengthen that finding in a crucial way. While age could reduce the information in ipsilateral motor cortex, it might also reduce information in contralateral motor cortex to a greater extent, such that ipsilateral cortex still provides compensatory (non-redundant) information. This question of redundant information can only be tested by combining voxels across hemispheres, as enabled by MVB. Indeed, the voxel weight spread measure from MVB in Experiment 1 hinted that older age might be associated with increased multivariate spread across hemispheres (Figure 4B left). Considered in isolation, this might support a compensatory role of ipsilateral motor cortex (contrary to Carp et al., 2011). However, MVB model comparison showed that adding these voxels did not lead to an age-related improvement in action decoding (i.e., this information was redundant to task performance, because it was already represented by the contralateral hemisphere). This illustrates the unique strength of an MVB approach, going beyond MVPA.

If the HAROLD pattern does not reflect compensation, what does the age-related hyper-activation of ipsilateral sensorimotor cortex reflect? One possible explanation is neural inefficiency, where older adults simply require greater neural and/or haemodynamic activity for the same computation (for review, see Barulli & Stern, 2013). Alternatively, there is growing evidence of neural dedifferentiation, whereby the functional specificity of brain regions reduces with age, such that additional areas (e.g., in the case of HAROLD, those that are ipsilateral) become involved in tasks that were not required when younger (for review, see Koen et al., 2020). Related to both ideas is the notion of task difficulty, illustrated by studies showing that younger adults activate similar areas to older adults, but only for higher demands (Reuter-Lorenz & Cappell, 2008). Task difficulty indeed influences ipsilateral motor cortex activity differently with age (e.g., Seidler et al., 2004; Verstynen et al., 2005). The fact that we observed the inverse age effect during the boost analysis in Experiment 1 (i.e., a simple detection task) but not Experiment 2 (i.e., a more demanding, decision-making task) might be relevant, but this remains purely speculative because the difference could simply be attributed to power, given that Experiment 2’s sample was an order of magnitude smaller.

Another non-compensatory account of HAROLD is motor disinhibition. Transcranial Magnetic Stimulation (TMS) approaches have shown that movement-related motor cortex activity inhibits ipsilateral motor areas (Schambra et al., 2003; Lee et al., 2003; Sohn et al., 2003; Kobayashi et al., 2004; Vercauteren et al., 2008) and, crucially, that these mechanisms attenuate (Peinemann et al., 2001), or even reverse (Rowe et al., 2006; Talelli et al., 2008b), with age. In other words, increased ipsilateral activation could be the result of reduced interhemispheric/transcallosal inhibition (Ferbert et al., 1992; Lee et al., 2003; Plewnia et al., 2003; Naccarato et al., 2006; Talelli et al., 2008a; Langan et al., 2010; McGregor et al., 2011; Wang et al., 2016; Burianová et al., 2020). This is consistent with age-related disruption of corpus callosum integrity (e.g., Ota et al., 2006; Giorgio et al., 2010; Langan et al., 2010; also see Lenzi et al., 2007; Cox et al., 2015) and of functional connectivity between left and right motor cortices (Langan et al., 2010), as well as their concentrations of glutamate (Kaiser et al., 2005). Comparable inhibitory mechanisms have been proposed for memory (e.g., Logan et al., 2002; Gazzaley et., 2005; de Chastelaine et al., 2011) and, for motor control, this provides plausible explanations of why older adults commit unintended mirror movements more often than younger adults (e.g., Koerte et al., 2010). This hypothesis could be examined by testing inter-hemispheric structural and functional connectivity in samples like Cam-CAN.

Finally, note that our results are limited to finger key-presses (whether simple or choice), and it is possible that better evidence for compensation within the HAROLD framework would come from more complex motor tasks (e.g., grasping; Knights et al., 2021). Another important limitation to consider (which also applies to, but is rarely addressed by, prior studies) is the degree to which age-related effects could be driven by increased noise in the fMRI data, for example due to greater (uncorrectable) head motion (e.g., Geerligs et al., 2017) or age-related changes in neurovascular coupling (e.g., D’Esposito, et al., 2003). While the simple explanation that some of our results are driven by noisier data in older adults might weaken the classical power to detect significant age effects, this would not explain the high Bayes Factors we found for the null hypothesis that age does not interact with ipsilateral activation in predicting behaviour (Figure 4B). Likewise, if estimates were noisier in older adults, then successful decoding should have been less common for these participants, yet successful decoding was found for almost every participant in Experiment 1, while Experiment 2 showed the opposite pattern, where the likelihood of successful decoding increased with age. It is possible that the age effects we found in ipsilateral sensorimotor cortex were purely vascular (e.g., owing to a weaker neurovascular coupling, a form of the “inefficiency” hypothesis discussed above), rather than neural. However, when adjusting task-activations for resting-state fluctuation amplitudes (RSFA), which are assumed to capture vascular reactivity, Tsvetanov et al. (2015) found that the increase of ipsilateral motor cortex with age in the same Cam-CAN data used for Experiment 1 was one of few age-related effects to survive adjustment, suggesting it is not solely a vascular effect. Another limitation of the present study is that the sample was cross-sectional, which limits inferences to individual differences in birth year (and associated potential generational differences), rather than about the specific longitudinal changes that occur with age (e.g., Anstey et al., 2003). Future longitudinal studies could address this.

In conclusion, our behavioural and multivariate approaches both contradicted the hypothesis that HAROLD is compensatory. Instead, results suggested that, at least in the case of ipsilateral motor cortex activity evoked by finger movements, this activation in older adults is non-specific, perhaps reflecting neural inefficiency or motor disinhibition.

## Acknowledgments

The Cambridge Centre for Ageing and Neuroscience (Cam-CAN) research was supported by the Biotechnology and Biological Sciences Research Council (Grant No. BB/H008217/1). The project has also received funding from the European Union’s Horizon 2020 research and innovation programme (‘LifeBrain’, Grant Agreement No. 732592), which supported E.K.; R.N.H. was supported by the Medical Research Council (SUAG/046 G101400); We thank Alex Quent for statistical advice.

